# Individual differences in behavioral flexibility predict future volitional ethanol consumption in mice

**DOI:** 10.1101/2021.09.21.460866

**Authors:** Ellen M Rodberg, Elena M Vazey

## Abstract

Cognitive control is key to regulating alcohol intake and preventing relapse. Behavioral inflexibility can prevent adaptive strategies such as mindfulness or other relapse-prevention behaviors. In a mouse model we investigated whether individual variability in behavioral flexibility (using attentional set-shifting task; ASST) predicts future alcohol intake. Adult male and female C57BL/6J mice were subjected to ASST using a bowl digging paradigm where mice identify a baited bowl based on compound odor and textural cues. This was completed prior to any alcohol exposure. Individual performance across mice varied within the group. We integrated several metrics, specifically ASST stage completed, trials to completion and errors performed to produce an individual performance index measure of behavioral flexibility. After, ASST mice were trained to drink ethanol (15%, v/v, 1hr/day) for 3-4 weeks until intake stabilized. Using this prospective approach, we identified an inverse relationship between behavioral flexibility and drinking - less flexible mice had a propensity to consume more alcohol. Similar relationships have been identified previously in non-human primates and rats. Our results show that the relationship between alcohol and cognitive flexibility is a robust trait that is conserved across species and can be used in mice to study neural substrates underlying these behaviors.

**HIGHLIGHTS:**

- ASST can be used to examine individual differences in cognitive function in mice
- Behavioral inflexibility is related to future higher alcohol consumption
- Executive function can be used as a predictive risk factor for alcohol intake
- This relationship in mice supports previous findings across species

## INTRODUCTION

Excessive alcohol use is associated with negative health, economic, and societal impacts (Global Burden of Disease Alcohol Drug Use, 2018). Identifying the risk factors and indices that are predictive of a propensity to consume increased levels of alcohol is important for prevention, treatment, and a deeper understanding of the biological underpinnings of excessive drinking behavior. Although major factors influence the risk of excessive alcohol use, complex interactions are present at the individual level. Individual differences in alcohol consumption are well documented in human, primate, and rodents’ studies (Spoelder et al., 2017; Vivian et al., 2001). Understanding individual level variables that have a strong predictive relationship with alcohol intake may identify those most at risk of excessive drinking and development of alcohol use disorder (AUD).

Regulation of alcohol intake at the individual level requires executive function, particularly behavioral flexibility, the ability to alter and update behavioral strategies according to a changing environmental context (Berg, 1948; Birrell & Brown, 2000; Brown & Tait, 2016; Young, Powell, Geyer, Jeste, & Risbrough, 2010). Decreased executive functioning, cognition, and behavioral flexibility has been frequently demonstrated in humans, primate, and rodents as a consequence of excessive alcohol use (Gass et al., 2014; Houston et al., 2014; Kroener et al., 2012; Rodberg et al., 2017; Shnitko, Gonzales, Newman, & Grant, 2020; Trick, Kempton, Williams, & Duka, 2014). Recently, an association between lower behavioral flexibility and the likelihood of future heavy drinking has been identified in non-human primates and rats (De Falco et al., 2021; Shnitko, Gonzales, & Grant, 2019) but it is not clear whether this prospective relationship is seen in other models of alcohol use. Behavioral flexibility as a potential predictive factor that is further impacted by excessive alcohol intake suggests an important bidirectional relationship with alcohol consumption.

In non-human primates, individual differences in performance on the attentional set shifting task (ASST) are predictive of future ethanol (EtOH) consumption, latency to drink, and drinking patterns (Shnitko et al., 2019). Here we aimed to identify whether behavioral flexibility in mice is predictive of individual differences in future EtOH consumption. Should the relationship identified in non-human primates back-translate across species we expect that mice who show poor ASST performance will show a greater escalation in their drinking over time and will exhibit higher average as well as single day consumption of EtOH.

## METHODS

### Animals

Adult C57/BL/6J male and female mice (n= 17, male n=9, female n=8; 8 weeks old) purchased from Jackson Labs (Bar Harbor, ME) were single housed on a reverse light cycle (11PM-11AM lights on). Animals had access to ad libitum chow and water before and after set shifting behavioral training and testing. During behavioral training and testing animals were given access to wet chow for 2hr/day after training/testing daily. Animal’s weight was monitored to ensure maintenance of 80% of baseline weight. All procedures were approved by the Institutional Animal Care and Use Committee at the University of Massachusetts at Amherst in accordance with the guidelines described in the U.S. National Institutes of Health Guide for the Care and Use of Laboratory Animals.

Outline of experimental timeline is illustrated in Figure 1A. Animals were handled daily by experimenter prior to habituation, training, and testing on an attentional set-shifting task. After behavioral experiments, animals’ EtOH drinking was obtained over 3-4 weeks.

**Figure 1.**
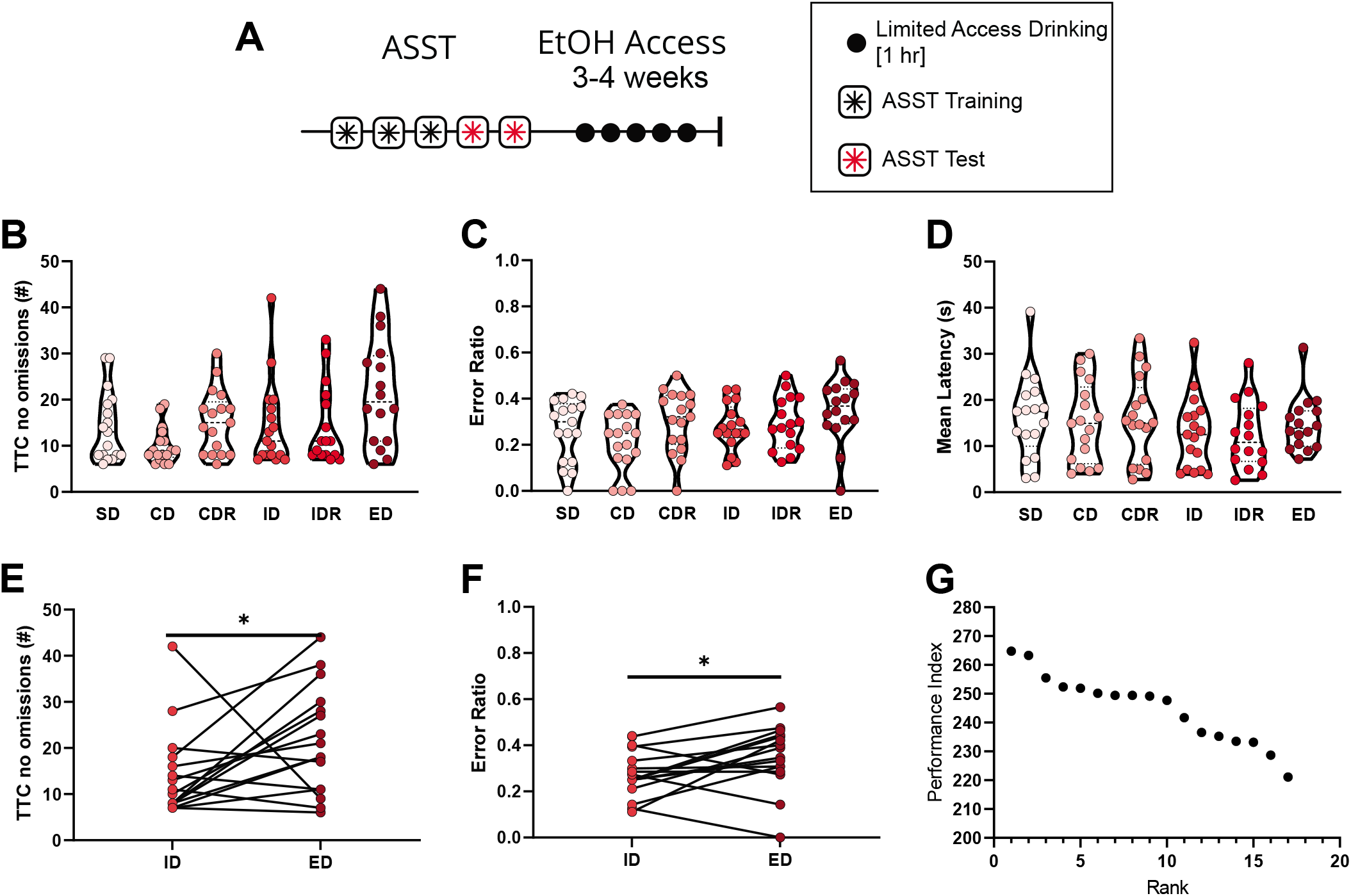
Performance on the attentional set shifting task as a basis for performance index. (**A**) Timeline of experiment. After 3 days of attentional set shifting habituation and training, animals were tested on the attentional set shifting task over 2 days. After the ASST, 1hr/day EtOH consumption was measured for 5 days a week over 3-4 weeks. (**B**) Trials to criterion (TTC), not including omitted trials, on each stage of the ASST. As subsequent stages became more difficult, the TTC increased (F (3.456, 53.91) = 3.405, p=0.0192; REML ANOVA). (**C**) Ratio of error trials out of all trials for each stage of the ASST. There was no effect of the stage on the ratio of errors to total trials per stage (F (3.709, 57.86) = 2.340, p=0.0702; REML ANOVA). (**D**) Average latency for an animal to respond during each stage of the ASST. As the staged became more difficult, there was no significant change in the average latency to make a response during the ASST (F (2.466, 38.47) = 1.019, p=0.3835; REML ANOVA). (**E**) The formation of an attentional set was validated by an increase in the TTC from the ID stage (14.41 ± 2.257) to the ED stage (21.50 ± 2.893) (p=0.0300; Wilcoxon test). (**F**) Ratio of incorrect trials to TTC on the ED stage (0.3494 ± 0.03414) compared to the ID stage (0.2767 ± 0.02436) was significantly increased (p=0.0302; Wilcoxon test). (**G**) Performance index (PI) was calculated for each animal by adding the normalized value of stage reached, average number of trials to criterion, and ratio of errors to trials. Higher PI values indicate a better performance on the ASST, and lower values indicate worse performance. Graph shows how the rank of animal performance (rank of 1 indicates the best performing animal) corresponds to the highest PI. Each dot represents the performance of one individual mouse.

### ASST training

ASST was performed as previously described (Rodberg et al., 2017) prior to any exposure to EtOH. Here, we used two stimuli dimensions (odor and texture). The ASST apparatus consisted of a custom-made acrylic box (49 x 20 x 21 cm) with a removable divider separating the waiting arena from two equal-sized choice compartments in the rear of the box. Animals were trained to dig for chocolate sucrose pellets (5 mg; Test Diet, Richmond, IN) in porcelain pots filled with scented digging media (5.5 x 5.5 x 3 cm) on top of a textured platform (10 x 7.5 x 2.5 cm) in the choice compartments. Crushed chocolate sucrose pellets were mixed into all scented digging media to mask the reward odor. Media were refilled on each trial, and pots were replaced after each stage and thoroughly cleaned after use. Each pot was designated to a specific odor to avoid cross-contamination of odor stimuli. Separate platforms of each texture were used for the same groups of mice to avoid introducing scents from unfamiliar mice. ASST arenas were thoroughly cleaned with 70% isopropyl between each animal. Twenty-four hours prior to and during ASST, mice were restricted to 2-hour daily access to wet food.

### Training day one

Training and habituation began three days prior to testing. The day before training, animals were food restricted to increase motivation for sucrose rewards. On day one of training animals were placed in an ASST arena for ten minutes to habituate to the testing arena. Animals were initially placed in the waiting area closed off by a divider. The divider was removed revealing the two choice compartments each with a ramekin containing sucrose reward pellets. Training concluded after animals ate sucrose pellets from each ramekin. To reduce neophobia, an empty ramekin with sucrose pellets were left in each animal’s homecage overnight.

### Training day two

Day two of training was similar to day one with the addition of unscented bedding sprinkled of top of the sucrose pellets in the ramekins. When animals retrieved the sucrose pellet from each ramekin, they were place back in the waiting area and ramekins were rebaited with sucrose with unscented bedding sprinkled on top. The divider was then removed, and animals could retrieve sucrose pellets from each ramekin. This process was repeated, adding more unscented bedding to the ramekin each time until animals would reliably dig from the ramekins to retrieve the sucrose reward. A ramekin with sucrose pellets buried in sawdust was left in the animals homecage overnight.

### Training day three

On day three of training, animals had to dig in bedding filled ramekins to receive a sucrose reward. Animals were exposed to and had to retrieve a sucrose pellet from all 6 textured platforms (fine sandpaper, coarse sandpaper, burlap, silk, tinfoil, velvet) and from all 6 scented beddings (cloves, cinnamon, basil, thyme, cumin, sage). Pairs of scented bedding and textured platforms were randomly paired and presented. Training concluded after animals received a sucrose pellet from each texture and scented bedding. A ramekin filled with unscented bedding and a sucrose pellet was placed in their homecage overnight.

### Testing

We performed a six-stage ASST consisting of simple discrimination (SD), compound discrimination (CD), compound discrimination reversal (CDR), intradimensional discrimination (ID), intradimensional discrimination reversal (IDR), and then extradimensional discrimination (ED). Testing took place over two days to prevent reward satiation with SD-CDR on day one and ID-ED on day two. The first day of training animals went through the first 3 stages of the ASST; SD, CD, and CDR. Animals were randomly assigned to protocols that started with either texture or scent, and the order of presented textures and scents were randomized. Each testing stage began with four discovery trials in which the animal could explore and dig in both choice compartments until they found the reward. Discovery trials were excluded from analysis. On test trials, the divider was closed and the response recorded when the animal approached the first digging pot. The divider prevented animals from entering more than one compartment per trial. Choices were recorded as correct, incorrect, or omissions (no choice after 60 seconds). Criterion was 6/8 consecutive correct choices, while failure to complete a stage was 20 omissions or 60 maximum trials. The rewarded choice compartment and exemplar pairing were randomized to be counterbalanced by side. Relevant stimulus dimension was balanced between animals and within groups (e.g., odor to texture).

### Drinking

Mice were given one-hour, one-bottle EtOH access (15%, v/v) beginning three hours after onset of the dark cycle. Drip was estimated with an EtOH bottle placed in an empty cage on the cage racks. Drip was subtracted from each animal’s EtOH consumption. Mice were weighed weekly and EtOH consumption was converted to g/kg. Data was analyzed using average consumption during each week.

### Performance Index

Performance indexes (PI) were calculated based off a modified version of a previously reported paper (Shnitko et al., 2017). The variables used to calculate PI were the stage reached in the ASST task, average number of trials (not including omissions) to complete each stage, and the average ratio of errors to total trials. To normalize each factor that contributed to the PI, the following equation was used:

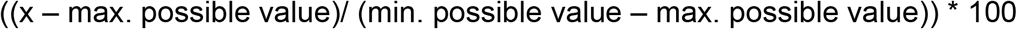

Performance index was calculated by adding the normalized value of stage reached, average number of trials to criterion, and ratio of errors to trials. Higher PI values indicate a better performance on the ASST, and lower values indicate worse performance.

### Statistics

For all analyses, p < 0.05 was used as the acceptable α level. Behavioral data was analyzed using GraphPad Prism v8 (San Diego, CA). Behavioral data is presented in violin plots with individual data points or mean ± SEM after confirming normality (Shapiro-Wilk) unless noted otherwise in the figure legend. Comparisons between groups were made with REML ANOVA and Sidak’s post hoc tests. Correlations were made using a one-tailed Spearman r. All graphs were composed using GraphPad Prism and compiled in Adobe Illustrator CS (San Jose, CA).

## RESULTS

### Attentional Set Shifting Task Performance

To investigate individual differences in behavioral flexibility prior to EtOH consumption, we measured animals’ using a 6-stage ASST. Training and test occurred one week prior to EtOH exposure (Fig 1A). There was an overall effect of ASST stage on the number of trials to complete each stage (excluding omissions) with increased trials needed as subsequent stages increased in difficulty (F (3.456, 53.91) = 3.405, p=0.0192; REML ANOVA; Fig. 1B). There was no effect of the stage on the ratio of errors to total trials per stage (F (3.709, 57.86) = 2.340, p=0.0702; REML ANOVA; Fig. 1C). There was also no effect of the stage on the average latency to complete a trial (F (2.466, 38.47) = 1.019, p=0.3835; REML ANOVA; Fig. 1D).

To specifically measure the formation of an attentional set during ASST we compared performance on extradimensional shifts (ED) to performance on the intradimensional shift stage (ID). The number of trials necessary to complete the ID and ED stage (not including omissions) was significantly increased in the ED (21.50 ± 2.89) stage compared to the ID (14.41 ± 2.26) stage (p=0.0300; Wilcoxon test; Fig. 1E). Furthermore, the ratio of incorrect to total trials was significantly increased in ED (0.3494 ± 0.034) compared to ID (0.2767 ± 0.024) (p=0.0302; Wilcoxon test; Fig. 1F).

To compare behavioral flexibility or cognitive performance across animals, performance index (PI) was calculated for each animal (modified from Shnitko et al., 2017). PI was calculated based on the stage reached, average trials (no omissions) to criterion, and ratio of incorrect to total trials. PIs ranged from 221.08 to 264.77 where animals performing poorly on ASST had lower PI than animals that performed well during the ASST (Fig. 1G).

### Volitional alcohol consumption and relationship to ASST

The average EtOH intake during one-hour sessions increased over the three-four weeks of EtOH access (week one (1.52 ± 0.12 g/kg), week two (2.06 ± 0.13 g/kg), week three (2.21 ± 0.13 g/kg), week four (2.39 ± 0.17 g/kg); Fig 2A). All mice showed EtOH consumption consistent with previous reports of drinking behavior seen in C57BL/6J mice (Becker & Lopez, 2004; den Hartog et al., 2020; Rodberg et al., 2017). Individual EtOH consumption stabilized by the last two weeks of drinking (last week (2.28 ± 0.14), second to last week (2.18 ± 0.14), ( t(16)=0.8275, p=0.4201; Paired t-test)). Comparisons of EtOH consumption across time by sex revealed a main effect of sex (F (1, 15) = 5.385, p = 0.0348; REML ANOVA), week (F (2.589, 31.93) = 14.30, p<0.0001; REML ANOVA), and an interaction (F (3, 37) = 5.013, p=0.0051; REML ANOVA) (Fig. 2B). Post-hoc analysis revealed that females (week one (1.45 ± 0.22 g/kg), week two (2.21 ± 0.19 g/kg; p=0.0034), week three (2.55 ± 0.125 g/kg; p=0.0017), week four (2.67 ± 0.15 g/kg; p=0.0258); Sidak’s multiple comparisons) but not males (week one (1.60 ± 0.12 g/kg), week two (1.92 ± 0.17 g/kg; p=0.2005), week three (1.91 ± 0.17 g/kg; p=0.2050), week four (1.82 ± 0.17 g/kg; p=0.6093); Sidak’s multiple comparisons) increased their drinking compared to the first week of baseline drinking on all following weeks. The increase in EtOH consumption in females is consistent with previous reports (Eriksson and Pikkarainen, 1968; den Hartog et al., 2020). Overall, across all animals average EtOH consumption increased from week one to week four from 1.53 ± 0.12g/kg to 2.28 ± 0.14 g/kg (p=0.0002; Wilcoxon test).

**Figure 2.**
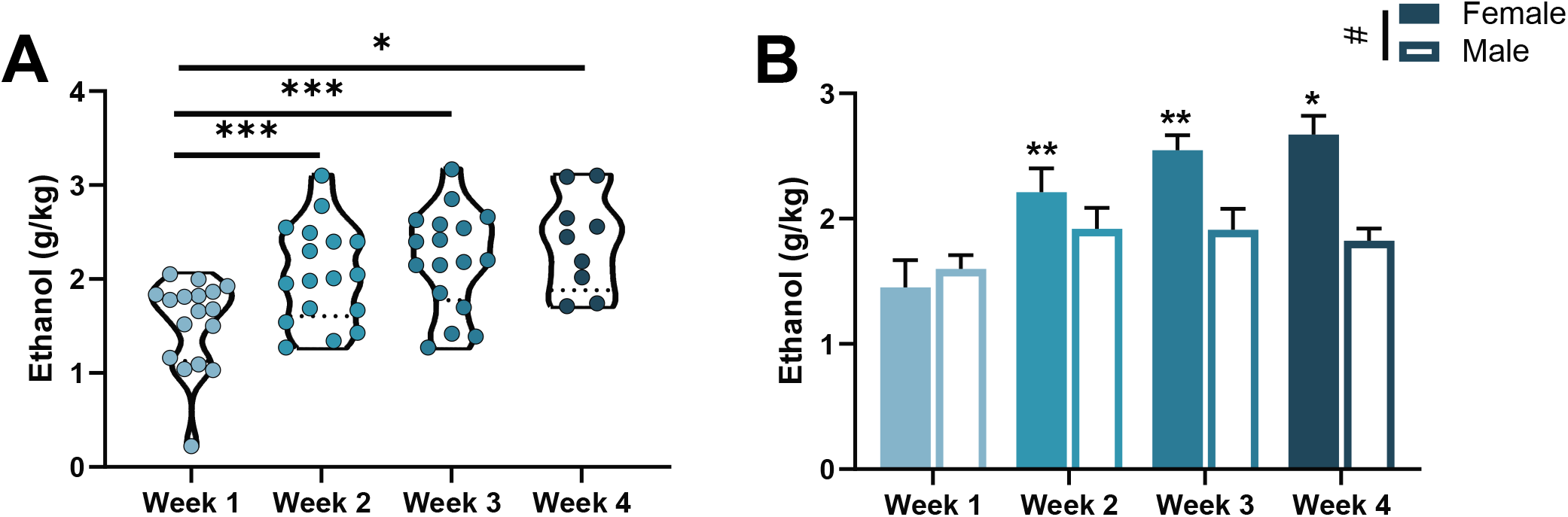
EtOH consumption escalates and stabilizes within a four-week period. (**A**) Average EtOH consumed in a 1hr period increased over the four-week period (F (2.045, 27.26) = 12.10, p=0.0002; REML ANOVA). Compared to week one (1.529 ± 0.1157 g/kg), EtOH consumption was significantly increased on week two (2.056 ± 0.1277 g/kg; p=0.0010), week three (2.209 ± 0.1302 g/kg; p=0.0010), and week four (2.389 ± 0.173 g/kg; p=0.0157). Each dot represents an animal’s average EtOH consumption, measured in g/kg, over a week. (**B**) Average EtOH consumption over the four-week period increased over time(F (2.589, 31.93) = 14.30, p<0.0001; REML ANOVA), was significantly different in males and females sex (F (1, 15) = 5.385, p = 0.0348; REML ANOVA), and there was a significant time and sex interaction (F (3, 37) = 5.013, p=0.0051; REML ANOVA). Females, but not males increased their EtOH consumption over each week compared to week one. (Females (week one (1.451 ± 0.2179 g/kg), week two (2.210 ± 0.1914 g/kg; p=0.0034), week three (2.546 ± 0.1212 g/kg; p=0.0017), week four (2.672 ± 0.1480 g/kg; p=0.0258); Sidak’s multiple comparisons). Males (week one (1.599 ± 0.1167 g/kg), week two (1.919 ± 0.1676 g/kg; p=0.2005), week three (1.910 ± 0.1698 g/kg; p=0.2050), week four (1.824 ± 0.1735 g/kg; p=0.6093); Sidak’s multiple comparisons).

To assess if performance during ASST was predictive of future EtOH consumption, we compared PI to the maximum daily EtOH consumption, EtOH escalation (average EtOH drinking on the last week compared to the first week), and the average EtOH consumption over the last two weeks of drinking. There was not a statistically significant correlation of PI and EtOH escalation (r=-0.2647, p=0.1517; Spearman r; Fig 3A) although there is a visible trend where lower values of PI had higher escalations. There was a significant negative correlation with PI and maximum EtOH consumption (r=-0.4255, p=0.0448; Spearman r; Fig 3B). Animals with lower PI, who performed worse in the ASST, had higher maximum one-day EtOH consumption. Furthermore, there was a significant negative correlation with PI and the average EtOH consumption over the last two weeks of drinking (r=-0.4632, p=0.0315; Spearman r; Fig 3C). This data suggests that animals with lower performance on the ASST, consumed larger quantities of EtOH on average and during individual drinking sessions.

**Figure 3.**
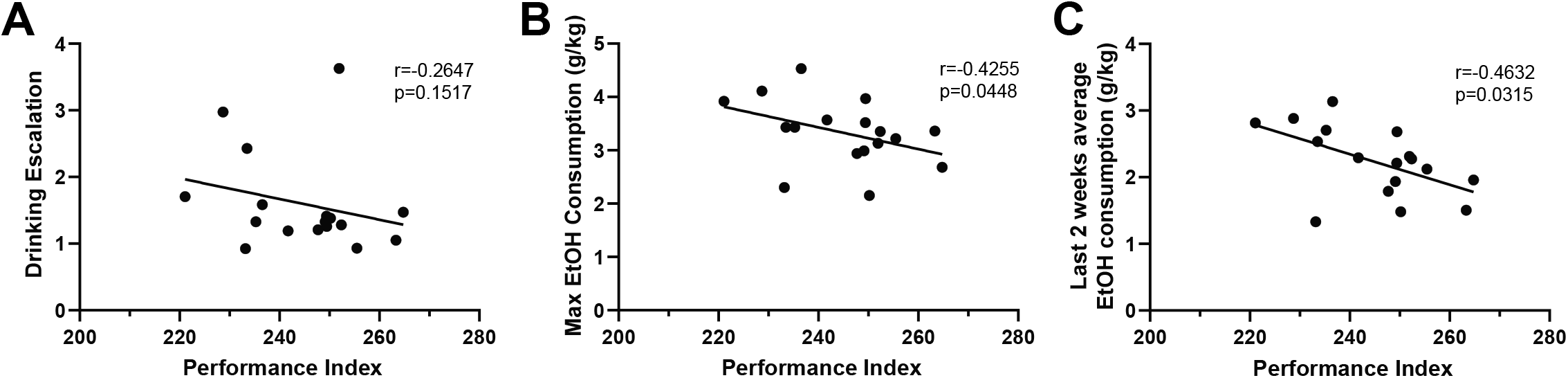
Lower behavioral flexibility is associated with increased EtOH consumption. (**A**) Correlation between PI and EtOH escalation (measured by average EtOH drinking on the last week compared to average EtOH drinking on the first week) was not statistically significant (r=-0.2647, p=0.1517; Spearman r). (**B**) Correlation between PI and the maximum EtOH consumption in one day of drinking was statistically significant (r=-0.4255, p=0.0448; Spearman r) where animals with lower PI had a higher maximum one-day EtOH consumption. (**C**) Correlation between PI and the average EtOH consumption over the last two weeks of drinking was statistically significant (r=-0.4632, p=0.0315; Spearman r) where animals with lower PI had a higher average EtOH consumption.

## DISCUSSION

In this study, we examined the relationship between behavioral flexibility measured in an ASST and propensity to voluntarily consume EtOH over three to four weeks. We identified that individual differences in measures of cognitive performance correlated with voluntary EtOH consumption. We found a significant inverse correlation in behavioral flexibility and both maximum EtOH consumption and average EtOH consumption as well as a trend in drinking escalation over three to four weeks.

Our data show cognitive performance was an effective predictive measure of future patterns of volitional EtOH consumption in mice. This finding is consistent with previous studies that found nonhuman primates with lower ASST performance were more likely to be later classified as high drinkers (Baker et al., 2017; Shnitko et al., 2019). Importantly, our data validates the correlation between cognition and future EtOH consumption in mice, an animal model often used in neurobiological studies of EtOH consumption. Furthermore, our results highlight individual differences in rodent baseline cognitive ability and propensity to consume EtOH are correlated without/prior to experimental treatments.

We identify that behavioral flexibility in mice is a feasible and useful metric for assessing risk for potential future alcohol consumption escalation. Excessive consumption of alcohol and AUD are characterized by increased compulsion to drink, explained by increased habitual behavior and an inability to control and inhibit behavioral responses. Resistance to change behavior in light of changing environmental conditions is an underlying component to both excessive alcohol consumption and a decreased behavioral flexibility. This supports previous findings across species that repetitive habitual behavior is associated with increased EtOH consumption in primates and alcohol preferring rats (De Falco et al., 2021; Shnitko et al., 2019). Evidence of this relationship in a rat strain specifically bred for high alcohol preference supports a biological basis underlying these co-occurring traits with potentially shared genetic and/or neural origins.

Our results add to the literature that there is a bidirectional relationship between cognitive function and increased alcohol consumption. It has been reported in humans and animal models that chronic alcohol consumption induces cognitive impairments in executive functioning (Bernardin, Maheut-Bosser, & Paille, 2014; Houston et al., 2014; Rodberg et al., 2017; Trick et al., 2014). This study, and others, have shown that decreased baseline cognitive ability is associated with higher levels of individual alcohol consumption (De Falco et al., 2021; Shnitko et al., 2019). Performance on ASST can be used as a readout of cognitive disruption to both predict proclivity to consume large amounts of alcohol and to measure cognitive deficits caused by alcohol consumption. Our results add to our understanding of behavioral inflexibility as a risk factor vs. a consequence of alcohol consumption, showing that cognitive deficits are not solely a result of alcohol consumption but that cognitive capacity increases individual vulnerability to high drinking phenotypes in a feed forward cycle.

Overall, we found that individual performance on an ASST task measuring behavioral flexibility was associated with future volitional EtOH consumption and ASST may be used as a predictive measure of the propensity or drive to consume increased quantities of alcohol in mice. We validated behavioral flexibility as a risk factor for alcohol intake in a laboratory setting, limiting confounding factors often present in human studies. Despite differences in experimental design and species, our findings are similar to results in primates and rats, suggesting that correlations between behavioral flexibility and EtOH consumption form a robust and highly translatable trait. There has been extensive research into the neurobiological substrates of executive function. Our findings provide evidence to support future studies on how neural systems underlying executive function may influence alcohol consumption across species to identify predictive biomarkers and phenotypes at risk for excessive alcohol use.

## FUNDING AND ACKNOWLEDGEMENTS

This work was supported National Institute of Health research grant U01AA025481. The authors would like to thank M Grampetro, M Nash, G Jacobs, R Davison, and C den Hartog for technical assistance with mouse drinking.

## AUTHOUR CONTRIBUTIONS

**Ellen Rodberg**: Methodology, Investigation, Analysis, Writing and Visualization. **Elena Vazey**: Conceptualization, Investigation, Writing – review and editing, Project administration, Supervision, Funding.

